# GO-Term Enrichment of Proteome-Scale Docking Profiles as a Biological Search-Space Reduction Layer for Protein Target Discovery

**DOI:** 10.64898/2026.09.15.751870

**Authors:** Chase Harms, Judith Klein-Seetharaman

## Abstract

Identifying protein targets from phenotype-first or mechanism-uncertain compounds remains difficult because proteome-scale docking can generate thousands of structurally plausible interactions per compound. We developed a workflow that converts ranked proteome-scale docking profiles into stable Gene Ontology (GO) Biological Process enrichment signatures and evaluates whether those signatures can reduce the candidate target space while preferentially retaining known drug-target relationships.

Docking targets were retained at the PDB-chain level, mapped to unique human gene identities, and analyzed with PANTHER overrepresentation against the screened structural gene universe. GO enrichment was evaluated from the top 25 through 505 ranked proteins in increments of 10, and a stable compound-level GO profile was selected using a Jaccard stability threshold of 0.80 across three consecutive transitions. Benchmarking used Yamanishi drug-target interactions, with 682 mapped compounds assigned to a prespecified development/held-out split (552/130) and evaluated at canonical, mechanistic, and fine-mechanistic biological resolutions.

In the full corrected benchmark, 642 compounds with usable canonical GO profiles showed greater within-class than between-class similarity (0.1019 versus 0.0931; delta = 0.0088; 100,000-permutation p = 0.00222), and canonical class explained 1.34% of multivariate GO-profile variation by PERMANOVA (p < 1e-4). Fine-mechanistic labels showed stronger organization in the full dataset (delta = 0.0245; PERMANOVA R-squared = 0.0976; both p < 1e-4). Held-out validation was more modest and metric-dependent: mechanistic labels were significant by PERMANOVA (R-squared = 0.0626, p = 0.0437), whereas the frozen fine-mechanistic analysis showed greater within-class similarity (0.1350 versus 0.1110; p = 0.038) and significant nearest-neighbor recovery (p = 0.0495), but not significant PERMANOVA (p = 0.119).

The principal held-out search-space experiment evaluated 107 compounds, 753,492 candidate protein rows, and 424 represented gold-standard targets. A direct GO gate retained 1.26% of candidates while retaining 11.32% of known targets (8.96-fold enrichment); ontology-propagated GO associations retained 5.10% of candidates and 21.46% of known targets (4.21-fold enrichment). At matched candidate-space sizes, GO-associated prioritization retained 27.59% versus 22.41% of known targets at approximately 5% of candidates, 39.39% versus 36.08% at 10%, and 58.73% versus 56.13% at 20%. By contrast, additive protein-level GO reranking was heterogeneous: among 605 evaluable compounds, 21.7% improved their best known-target rank, but mean reciprocal rank decreased from 0.0276 to 0.0189. These results support GO enrichment as an intermediate biological search-space reduction and prioritization layer rather than a universal direct target-scoring function.

## 1. Introduction

Phenotypic screening and observations of biological activity frequently identify compounds with useful or unexpected effects before the protein mechanisms responsible for those effects are fully understood. Determining the molecular targets responsible for such phenotypes is therefore a central problem in chemical biology, pharmacology, drug repurposing, and natural-product discovery [1].

Molecular docking provides one route for generating target hypotheses. Conventionally, docking is performed against a small number of proteins selected using prior biological knowledge. This is useful when a mechanism is already suspected, but it limits discovery of unexpected targets. The inverse strategy screens a compound against thousands of available human protein structures to generate a predicted structural targetome [2,3]. Such proteome-scale docking preserves a broad hypothesis space but creates a second problem: thousands of predicted interactions cannot reasonably be investigated experimentally, and docking-score differences alone are insufficient to establish physiological target identity [3].

We hypothesized that collective biological organization within a proteome-scale docking profile could provide an intermediate layer between raw structural ranking and individual target hypotheses. If high-ranking docking candidates are disproportionately associated with a coherent biological process, that process-level signal may be detectable even when no single docking score uniquely identifies the true target.

GO enrichment is suited to detecting such collective signals [7,8], but its application to docking data introduces several methodological questions: the appropriate depth of the docking list is not known a priori, enrichment can change as progressively weaker docking candidates are included, and enrichment against the entire human proteome can be biased when the actual structural screen covers a non-random subset of proteins [9,10].

We therefore developed a workflow that repeatedly evaluates GO Biological Process enrichment across progressively larger docking subsets, uses the screened structural gene universe as the enrichment background, and selects a stable enrichment depth based on reproducibility of top GO terms across consecutive cutoffs. We then distinguish two possible uses of the resulting biological information:

- Direct protein reranking: assign an individual candidate protein a higher score when its GO annotations overlap the compound-specific enriched GO processes.
- Biological search-space reduction: use enriched biological processes to define a smaller, biologically coherent region of the docking targetome for subsequent structural and experimental analysis.

The present benchmark tests both interpretations at increasing biological label resolution and, critically, on a prespecified held-out compound set. The resulting data support GO enrichment as a useful search-space reduction and contextualization layer, while arguing against its naive use as a universal additive protein-target score.

## 2. Materials and Methods

### 2.1 Human structural target library

A structural library of human proteins was assembled from experimentally determined structures deposited in the Protein Data Bank (PDB) [4]. Structures were filtered to Homo sapiens, resolution less than 2.5 angstroms, and sequence-identity grouping at 50%. Crystallographic waters, bound ligands, cofactors, and ions were removed according to the screening protocol. Structures were protonated assuming pH 7 and hydrogens were added. Each docking target was retained at the PDB-chain level to permit chain-specific mapping to the corresponding human gene. A total of 7,518 structures were gathered.

### 2.2 Proteome-scale molecular docking

Each compound was docked independently against the structural target library using AutoDock Vina [5]. Docking was performed as a blind docking screen, and the best docking score for each protein-compound pair was retained to construct a ranked structural targetome for each compound. [AUTHOR TO COMPLETE BEFORE SUBMISSION: AutoDock Vina version, docking-box definition, exhaustiveness, number of poses, and any scoring/preparation parameters not described above.]

### 2.3 Strict PDB-chain to target-identity mapping

Docking targets were mapped from PDB structure and chain identifiers to UniProt records [4,6] and subsequently to human gene identities. Chain-specific mappings were preferred and cached for consistent reuse. PDB-level mappings were used as a fallback only when the PDB mapped unambiguously to a single gene. Multi-gene PDB-level mappings were not propagated to every chain; instead, unresolved chains were resolved at chain level. For target-recovery analyses, candidate target identities were required to be unique at the gene level (or, when no gene identity was available, at a unique UniProt level). Rows with ambiguous multi-gene or multi-UniProt identity were excluded rather than allowing co-complex proteins to share aliases. This strict mapping procedure prevented a protein in a multicomponent structure from inheriting the identities or GO annotations of neighboring chains.

### 2.4 Construction of the GO enrichment background

Because the experimentally determined structural proteome is a non-random subset of the human proteome, the enrichment reference population was defined from genes represented in the docking target universe rather than from all human genes [9,10]. The resulting one-gene-per-entry background was supplied to PANTHER as the reference input list. This design tests whether a biological process is overrepresented relative to the proteins that could actually have been identified by the structural screen.

### 2.5 GO Biological Process enrichment and stability selection

GO enrichment was performed using the PANTHER overrepresentation framework [7,8] for Homo sapiens with GO Biological Process annotations, Fisher’s exact test, and false-discovery-rate correction [8,14]. Terms were retained at nominal p < 0.05 and FDR < 0.05 and ranked by fold enrichment. Enrichment was evaluated from N = 25 through N = 505 top-ranked proteins in increments of 10, retaining the top 25 enriched GO terms at each depth. Similarity between consecutive top-term sets was quantified using the Jaccard coefficient J(A,B) = |A intersect B| / |A union B|. A stable region was defined by J >= 0.80 across three consecutive transitions, and the first qualifying depth was selected as the compound-level GO profile.

### 2.6 Yamanishi benchmark and prespecified split

Known drug-target relationships were obtained from the Yamanishi enzyme, GPCR, ion-channel, and nuclear-receptor interaction datasets [11]. The combined reference contained 5,127 interaction records spanning 791 Yamanishi drug identifiers and 989 target identifiers. Compound structures in the docking benchmark were mapped to Yamanishi drug identifiers. After compound-resolution review and conservative mechanistic cleanup, 682 compounds were assigned to a prespecified split containing 552 development compounds and 130 held-out compounds. Group labels were never used during docking, GO enrichment, or stability selection.

### 2.7 Biological label resolutions

Three complementary biological label resolutions were evaluated. Canonical labels used the four Yamanishi target classes (Enzyme, GPCR, IonChannel, NuclearReceptor). Mechanistic labels subdivided compounds using known target genes into ArachidonicAcid, DNA, GPCR, NuclearReceptor, OtherEnzyme, OtherIonChannel, and Voltage-gated groups, while multi-mechanism or unclassified compounds were excluded from single-label class tests. Fine-mechanistic labels further subdivided these parent mechanisms by target family, such as monoamine metabolism, metalloproteases, carbonic anhydrases, adrenergic GPCRs, histamine GPCRs, lipid/prostanoid GPCRs, sex-steroid nuclear receptors, and voltage-gated sodium channels. Fine labels were derived only from known Yamanishi target identities and not from docking or GO results.

For the primary fine-resolution held-out analysis, class eligibility was frozen from development data before held-out GO-profile testing. Specific fine groups required at least 10 development compounds; broad catch-all groups were excluded from the primary fine test; held-out groups required at least two usable compounds so that within-group similarity was defined. This yielded 15 evaluable fine-mechanistic held-out groups comprising 58 compounds with usable GO profiles.

### 2.8 Compound-level GO-profile similarity

Each compound was represented by a binary vector indicating presence or absence of each selected GO term. Pairwise similarity was calculated using Jaccard similarity. Mean within-group similarity was compared with mean between-group similarity. Because pairwise similarities sharing compounds are not statistically independent, significance was evaluated by permuting group labels while preserving group sizes. Each reported class-level analysis used 100,000 permutations.

### 2.9 PERMANOVA and nearest-neighbor recovery

One-way PERMANOVA [12] was applied to the complete Jaccard distance matrix to test multivariate association between biological group and GO-profile position. The fraction of compounds whose nearest GO-profile neighbor belonged to the same biological group was also compared with a group-label permutation null distribution. Source-specific and pairwise tests were corrected using the Benjamini-Hochberg false-discovery-rate procedure [14].

### 2.10 Direct GO-assisted target reranking control

For each candidate target, GO support was computed from overlap between the candidate’s own direct or ontology-propagated GO Biological Process annotations and the compound’s selected enriched GO terms. Candidate identity and GO support were restricted to the candidate’s unique mapped target identity; co-complex genes could not contribute aliases or GO annotations. A fixed prespecified GO contribution of 0.20 was added to the normalized docking rank score. This weight was not optimized against target-recovery performance. Known-target recovery was summarized using best-known-target rank, mean reciprocal rank (MRR), Hit@K, Recall@K, and the fraction of compounds whose best known-target rank improved.

### 2.11 Held-out biological search-space reduction validation

The primary applied validation asked whether GO information could reduce the candidate target universe while preferentially retaining known interactions. This analysis was restricted to the prespecified held-out split and included compounds for which at least one known Yamanishi target was represented in the structural candidate universe. Four prioritization strategies were evaluated across candidate-space fractions: docking only; GO-associated candidates first; a GO-weighted ordering; and random filtering as a null expectation. Natural GO gates were also evaluated using direct annotation overlap and ontology-ancestor propagation. A GO-plus-docking safety-net strategy retained the GO-associated set together with top-ranking docking outliers. The main endpoint was known-target retention as a function of the fraction of candidate proteins retained.

### 2.12 Interpretation and robustness

All held-out analyses were interpreted independently of development-set exploration. Fine-mechanistic all-data analyses were treated as exploratory because group-level patterns had access to development compounds. Significance at one biological resolution was not interpreted as proof of categorical separation at another resolution. PERMANOVA was interpreted together with within-versus-between similarity and nearest-neighbor recovery because these statistics capture different aspects of multivariate structure. A formal multivariate-dispersion (PERMDISP) analysis remains a recommended robustness check before final submission because unequal within-group dispersion can influence PERMANOVA, particularly in unbalanced held-out groups.

## 3. Results

### 3.1 Benchmark coverage and corrected target-identity mapping

The benchmark contained 682 mapped compounds in the prespecified 552/130 development/held-out split. Not every compound contributed a usable profile to every downstream analysis because class eligibility, GO-profile availability, and known-target representation differed by endpoint. The corrected canonical all-compound profile analysis included 642 compounds, the mechanistic analysis included 638 compounds, and the exploratory fine-mechanistic analysis included 522 compounds. Held-out canonical and mechanistic profile analyses each included 116 compounds. The frozen fine-mechanistic held-out analysis included 58 compounds distributed across 15 development-qualified fine groups.

For target-level analyses, strict candidate-identity mapping excluded ambiguous multi-gene, multi-UniProt, and unmapped structural rows instead of merging co-complex identities. This correction removed the possibility that a frequently occurring complex component could inherit unrelated receptor or enzyme identities and artificially count as multiple known-target matches.

### 3.2 Biological information was detectable in GO profiles and increased at finer mechanistic resolution

Across 642 compounds with usable canonical profiles and 1,020 unique selected GO terms, mean within-class Jaccard similarity was 0.1019 compared with 0.0931 between classes (delta = 0.0088, p = 0.00222). Canonical class was associated with the complete GO-profile distance matrix by PERMANOVA (pseudo-F = 2.8960, R-squared = 0.0134, p < 1e-4), and 43.93% of compounds had a nearest GO-profile neighbor from the same canonical class (p < 1e-4).

At mechanistic resolution, 638 compounds across seven groups had mean within-group similarity of 0.1011 versus 0.0944 between groups (delta = 0.0067, p = 0.0157). PERMANOVA remained highly significant (R-squared = 0.0195, p < 1e-4), and nearest-neighbor source recovery was significant (30.88%, p = 6e-5).

The strongest all-data organization occurred at fine-mechanistic resolution. Among 522 compounds in 25 fine groups, within-group similarity was 0.1257 versus 0.1012 between groups (delta = 0.0245, p < 1e-4). Fine-group identity explained 9.76% of multivariate variation (PERMANOVA p < 1e-4), and nearest-neighbor fine-group recovery was also greater than expected by permutation (p < 1e-4). Several individual fine groups showed large FDR-controlled within-versus-other effects, including metalloproteases (delta = 0.1079, FDR = 0.00025), monoamine metabolism (delta = 0.1012, FDR = 0.000375), lipid/prostanoid GPCRs (delta = 0.1177, FDR = 0.00835), adrenergic GPCRs (delta = 0.0352, FDR = 0.00835), histamine GPCRs (delta = 0.0588, FDR = 0.0182), sex-steroid nuclear receptors (delta = 0.0378, FDR = 0.0418), and voltage-gated sodium channels (delta = 0.0566, FDR = 0.00533). These all-data subgroup results are exploratory and were not treated as independent held-out confirmation.

**Table 1.** Resolution-specific GO-profile validation. The frozen fine held-out R-squared should be interpreted cautiously because 15 groups were represented by only 58 compounds; the associated pseudo-F was 1.1101 and the permutation p-value was 0.119.

| Resolution | Scope | N compounds | Within Jaccard | Between Jaccard | Delta | Similarity p | PERMANOVA R2 | PERMANOVA p |
| --- | --- | --- | --- | --- | --- | --- | --- | --- |
| Canonical | All | 642 | 0.1019 | 0.0931 | 0.0088 | 0.00222 | 0.0134 | <1e-4 |
| Mechanistic | All | 638 | 0.1011 | 0.0944 | 0.0067 | 0.0157 | 0.0195 | <1e-4 |
| Fine mechanistic | All | 522 | 0.1257 | 0.1012 | 0.0245 | <1e-4 | 0.0976 | <1e-4 |
| Canonical | Held-out | 116 | 0.1006 | 0.0943 | 0.0063 | 0.170 | 0.0332 | 0.0585 |
| Mechanistic | Held-out | 116 | 0.1002 | 0.0951 | 0.0051 | 0.234 | 0.0626 | 0.0437 |
| Frozen fine | Held-out | 58 | 0.1350 | 0.1110 | 0.0240 | 0.038 | 0.2655 | 0.119 |

**Figure 1.**
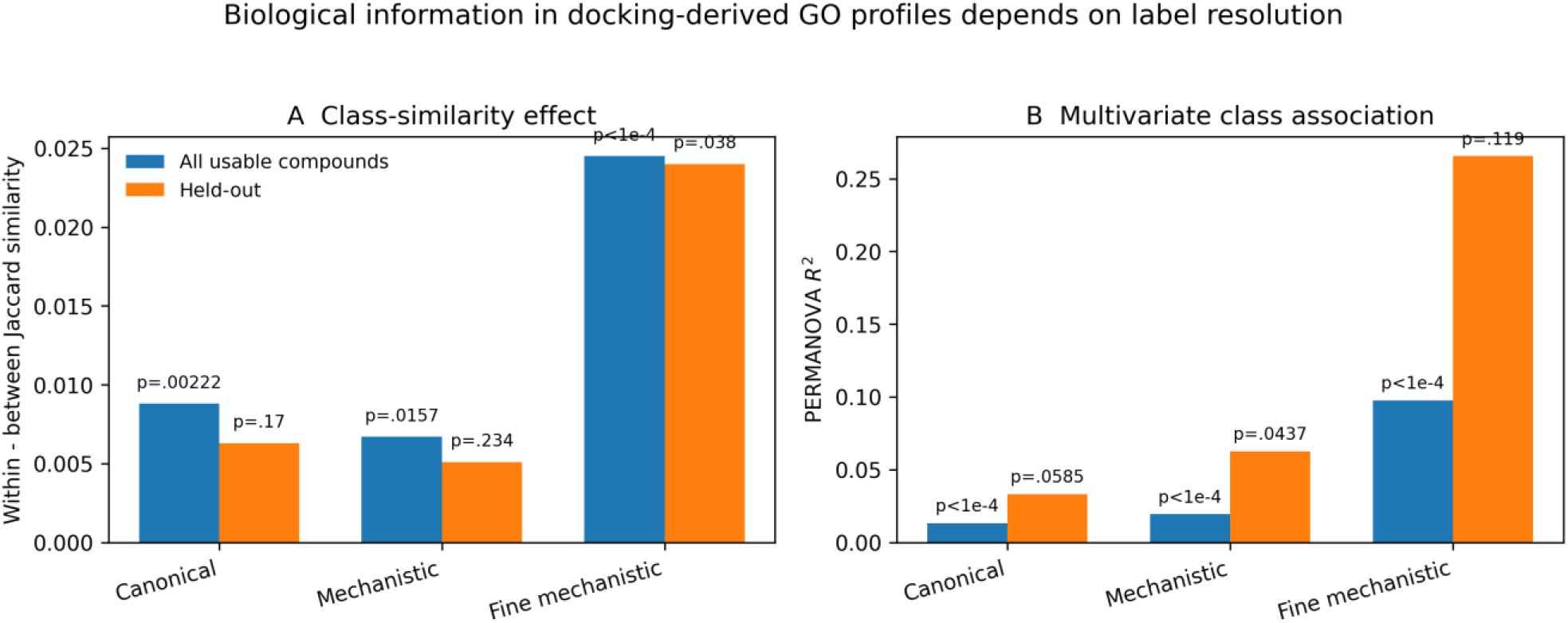
Biological signal across class resolutions. Panel A summarizes the within-minus-between Jaccard effect; Panel B summarizes PERMANOVA R-squared. All-data analyses show statistically detectable structure at each resolution, with the largest exploratory effect at fine resolution. Held-out evidence is mixed and metric-dependent: mechanistic PERMANOVA is significant, whereas the frozen fine analysis is significant by within-versus-between similarity but not by PERMANOVA. Fine held-out R-squared is therefore not interpreted as a validated 26.6% effect size.

### 3.3 Held-out validation supported partial, resolution-dependent generalization

The broad canonical held-out analysis did not meet conventional significance thresholds: within-class similarity was 0.1006 versus 0.0943 between classes (p = 0.170), PERMANOVA yielded R-squared = 0.0332 with p = 0.0585, and nearest-neighbor source recovery yielded p = 0.0688. Thus, broad Yamanishi class identity was not independently recovered with strong evidence in the held-out set.

At mechanistic resolution, the simple within-versus-between comparison also remained nonsignificant (0.1002 versus 0.0951; p = 0.234), but the complete multivariate distance structure showed a significant class association by PERMANOVA (R-squared = 0.0626, p = 0.0437). Nearest-neighbor recovery did not independently reach significance (p = 0.0821). The mechanistic held-out result therefore supports a modest multivariate class signal rather than uniformly tighter within-class clusters.

In the frozen fine-mechanistic held-out test, mean within-class similarity was 0.1350 versus 0.1110 between classes (delta = 0.0240, p = 0.038), and nearest-neighbor fine-group recovery was greater than expected by permutation (p = 0.0495). PERMANOVA was not significant (pseudo-F = 1.1101, p = 0.119). The held-out fine result therefore provides partial replication of fine-scale biological organization, but small group sizes (typically two to seven compounds) limit resolution-specific inference.

### 3.4 Direct additive GO reranking was heterogeneous and did not provide a universal target score

Target-level reranking was evaluable for 605 compounds. The median best-known-target rank improved numerically from 436 using docking alone to 409 after adding the fixed GO contribution, and 21.7% of compounds improved their best known-target rank. However, 73.9% worsened and only 4.5% were unchanged. Mean reciprocal rank decreased from 0.0276 to 0.0189, indicating that gains in a subset of compounds were offset by losses among compounds with highly ranked docking targets. In the held-out subset, 107 evaluable compounds showed the same qualitative pattern: median best-known-target rank shifted from 381 to 336, but mean reciprocal rank decreased from 0.0458 to 0.0177.

These results argue against treating additive GO overlap as a universal replacement for docking rank. The effect was strongly mechanism-dependent. In the corrected fine-group analysis, monoamine-metabolism compounds showed a median best-known-target rank shift from 73.5 to 12 across 20 compounds, with 55% improving. In the held-out monoamine subset, all three compounds improved and the median rank shifted from 105 to 10. Oxidoreductase compounds also showed frequent improvements, whereas several other biological families showed little or negative benefit. These subgroup observations support selective biological utility but are not a basis for a universal scoring rule.

**Figure 2.**
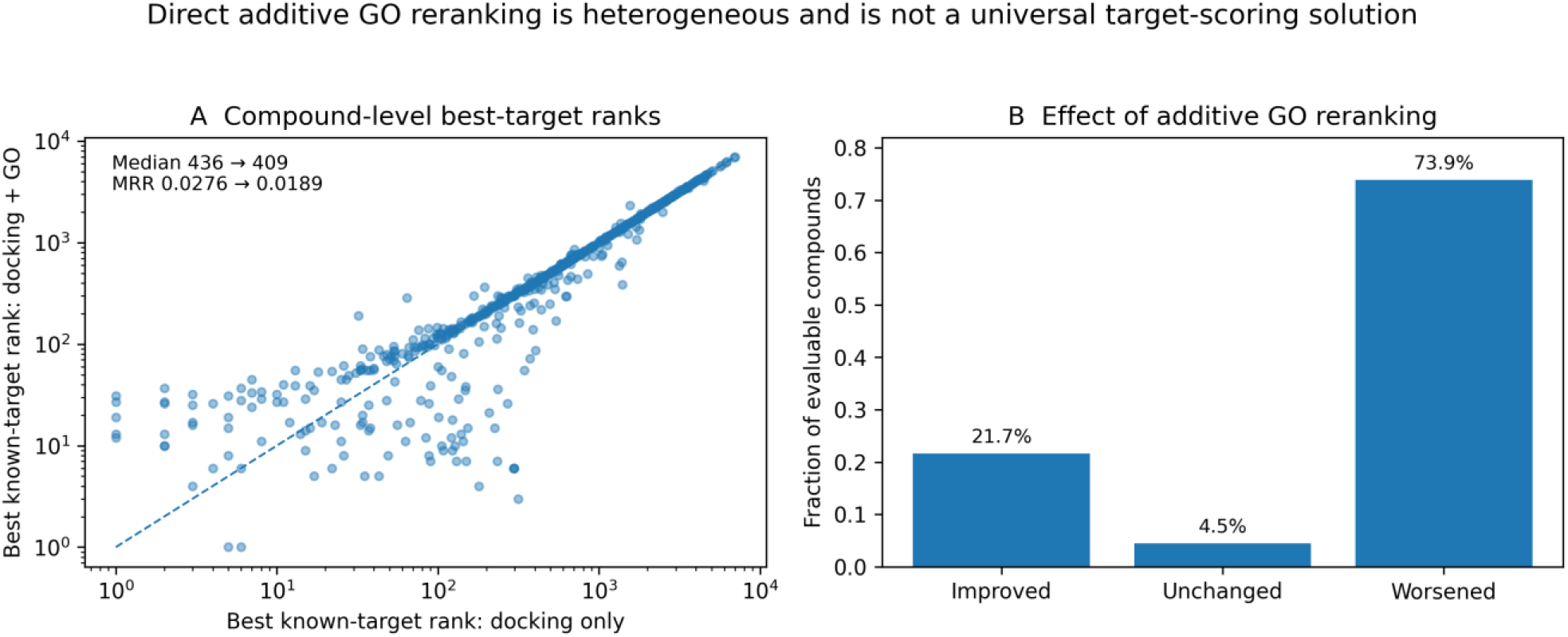
Corrected direct target-reranking control across 605 evaluable compounds. Panel A compares docking-only and additive GO-assisted best-known-target ranks on logarithmic axes; points below the diagonal improve. Panel B shows the fraction of compounds improved, unchanged, or worsened. The numerical median rank improves, but mean reciprocal rank declines because many already-high-ranking known targets are displaced.

### 3.5 GO prioritization improved held-out search-space efficiency

The principal search-space validation was performed exclusively on the prespecified held-out set and included 107 compounds, 753,492 candidate protein rows, and 424 represented gold-standard targets. A direct GO-annotation gate retained 9,521 candidate rows (1.26% of the screened candidate space) and 48 known targets (11.32%), corresponding to an 8.96-fold enrichment relative to candidate-space size. Allowing ontology-ancestor propagation retained 38,401 candidates (5.10%) and 91 known targets (21.46%), a 4.21-fold enrichment.

When candidate-space size was matched explicitly, GO-associated-first prioritization consistently exceeded the random-filter expectation and modestly exceeded docking alone at stringent and intermediate reductions. At approximately 5% of candidate proteins retained, GO-associated prioritization retained 27.59% of known targets compared with 22.41% for docking alone and 5.01% expected under random filtering. At 10%, target retention was 39.39% for GO-associated prioritization, 36.08% for docking alone, and 10.01% for random filtering. At 20%, the corresponding values were 58.73%, 56.13%, and 20.01%. By 50% of candidates retained, GO-associated and docking-only strategies converged at 87.50% target retention.

The same pattern was apparent when asking how much candidate space was required to reach fixed target-retention goals. GO-associated prioritization required approximately 15% of candidates to retain 50% of known targets, compared with 20% for docking alone. To retain 70% of known targets, GO-associated prioritization required approximately 30% of the candidate space, compared with 40% for docking alone. At 80% and higher retention goals, the two strategies converged.

**Table 2.** Held-out known-target retention at selected candidate-space sizes. Values for ∼1% and ∼2.5% are weighted across the 107 evaluable held-out compounds; the remaining selected values agree with the held-out search-space reduction report.

| Candidate fraction retained | GO-associated first | Docking only | Random expectation |
| --- | --- | --- | --- |
| ~1% | 11.79% | 6.13% | ~1% |
| ~2.5% | 20.28% | 12.74% | ~2.5% |
| ~5% | 27.59% | 22.41% | 5.01% |
| ~10% | 39.39% | 36.08% | 10.01% |
| ~20% | 58.73% | 56.13% | 20.01% |
| ~50% | 87.50% | 87.50% | 50.00% |

**Figure 3.**
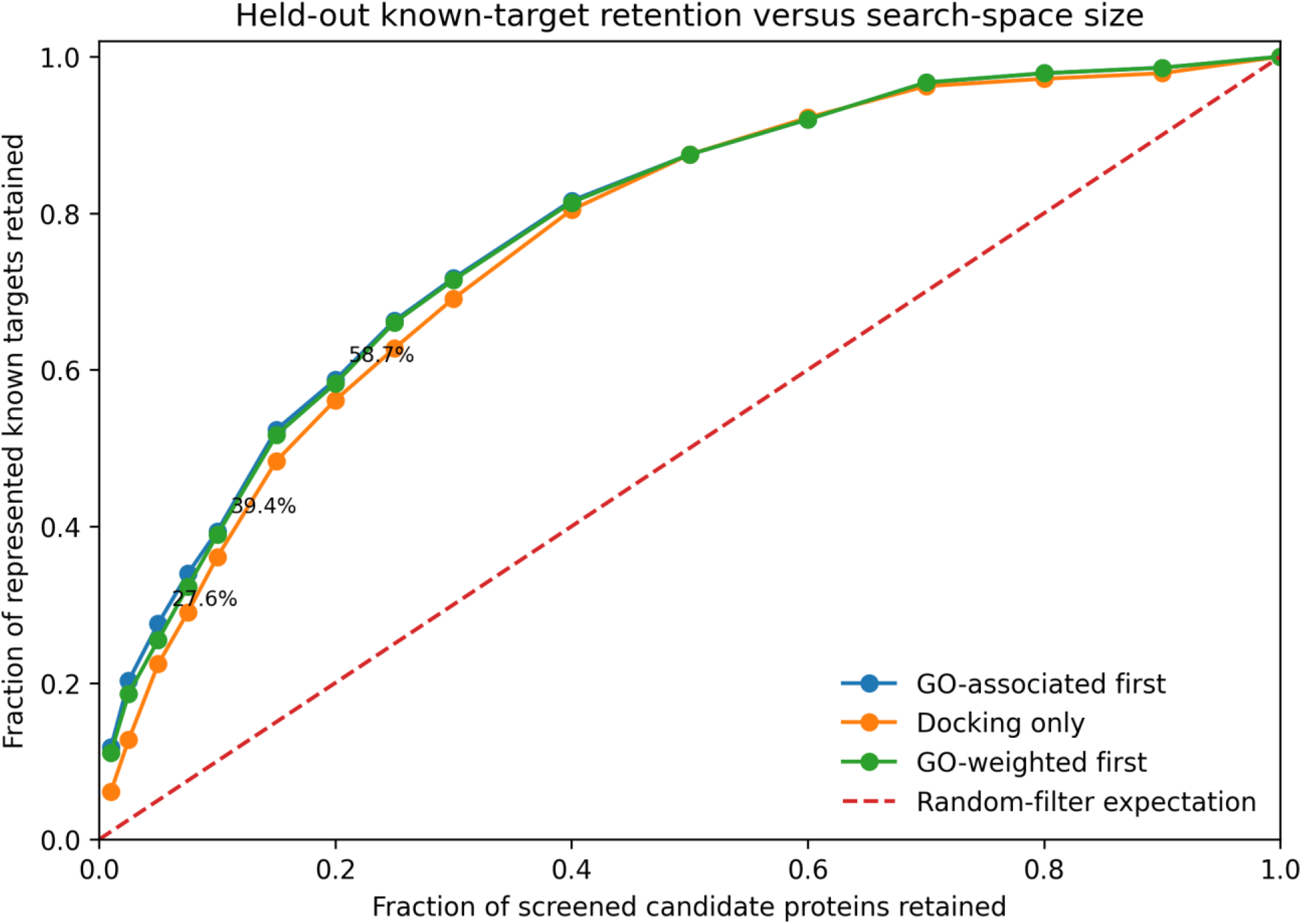
Held-out known-target retention versus candidate-space size. GO-associated-first and GO-weighted strategies are compared with docking-only prioritization and the random-filter expectation across 107 compounds and 424 represented known targets. The GO-associated strategy shows its largest advantage at stringent search-space sizes and converges with docking-only performance as progressively more of the targetome is retained.

### 3.6 Mechanism-specific examples illustrate where biological filtering can be especially effective

The search-space advantage was not uniform across mechanisms. In held-out monoamine-metabolism compounds, GO-associated-first prioritization retained all six represented known targets while retaining approximately 5% of the candidate space. For held-out oxidoreductase compounds, GO-associated-first retained all five represented targets by approximately 7.5% of candidate space, whereas docking alone retained only 20% at approximately 10% of candidates. Held-out phosphodiesterase compounds retained 57.1% of represented targets with GO-associated prioritization at approximately 10% of candidates versus 33.3% with docking alone. These subgroup sizes are small and therefore illustrative rather than definitive, but they are consistent with the broader conclusion that GO-based reduction is most useful in biological contexts that generate coherent process-level signatures.

## 4. Discussion

### 4.1 Proteome-scale docking contains recoverable biological information beyond individual docking scores

The corrected benchmark demonstrates that collective GO Biological Process structure can be recovered from proteome-scale docking profiles. This structure is statistically detectable in the full benchmark and becomes substantially stronger at finer mechanistic resolution, suggesting that docking-derived biological organization is more closely aligned with specific target families or mechanisms than with very broad pharmacological classes. At the same time, the held-out analyses show that this signal is not absolute: broad canonical classes do not reproduce strongly, mechanistic classes show modest multivariate held-out structure, and fine classes show partial replication that depends on the statistic used.

This pattern is biologically plausible. Compounds can interact with multiple proteins, target families can participate in overlapping pathways, and GO annotations capture downstream and shared biological processes rather than mutually exclusive mechanistic labels. A useful biological representation therefore need not produce sharply separated categorical clusters. The modest held-out effects are more consistent with a continuous, overlapping functional space than with a hard classifier.

### 4.2 Search-space reduction is the strongest validated application of the GO layer

The held-out search-space experiment provides the clearest practical validation of the method. Direct GO annotation identified a very small subset of candidate proteins that was strongly enriched for known targets, and ontology-propagated associations increased target coverage while retaining only a small fraction of the screened targetome. When search-space size was explicitly matched, GO-associated prioritization improved known-target retention over docking alone most clearly at stringent candidate fractions. This is precisely the regime in which a proteome-scale screen is difficult to use experimentally: thousands of plausible structures must be reduced to a manageable set for follow-up.

The practical role of GO enrichment is therefore not to declare a single molecular target. It is to identify biologically supported regions of a large structural targetome and to move those regions forward for more expensive target-level evaluation. The observed advantage over random filtering confirms enrichment of known biology, while the more modest advantage over docking alone indicates that GO contributes complementary rather than dominant information.

### 4.3 Biological process enrichment and individual target prediction are different inference problems

The direct reranking results clarify an important distinction. A compound-level GO profile can be biologically informative even when annotation overlap is not a reliable universal score for individual proteins. A molecular target can influence an enriched process without carrying the exact selected annotation, and many proteins annotated to an enriched process may not be direct binders. This hierarchy can be summarized as: compound -> molecular target -> pathway or network -> cellular response -> GO process. Information at the process level can therefore improve regional prioritization without monotonically improving every individual target rank.

The corrected analysis also shows that reranking benefit is heterogeneous rather than absent. Approximately one fifth of evaluable compounds improved their best known-target rank, and some mechanistic families showed substantial rescue. The simultaneous decline in mean reciprocal rank, however, demonstrates why these successes should not be generalized into a universal additive scoring model. A safer architecture is to use GO for candidate gating or prioritization while retaining high-affinity docking outliers as an orthogonal discovery tier.

### 4.4 Implications for phenotype-driven target discovery

The workflow is particularly suited to settings in which a biological phenotype is known but its initiating molecular target is uncertain. Proteome-scale docking preserves a broad structural hypothesis space; stable GO enrichment then identifies biological processes disproportionately represented among high-ranking candidates. These compound-derived processes can be compared with phenotype-associated biology from literature, omics, genetics, or experiments. Proteins at the intersection can be prioritized for pose inspection, binding-site analysis, higher-accuracy rescoring, molecular dynamics, expression and pathway analysis, biochemical assays, or perturbation studies.

This approach maintains discovery potential because the initial screen is not restricted to proteins already associated with the phenotype. At the same time, a two-tier downstream strategy prevents the GO layer from becoming a hard filter: biology-supported candidates can be investigated alongside exceptional docking outliers that fall outside the enriched process space.

### 4.5 Strict chain-level mapping is essential for target-fishing validation

A key methodological lesson from the benchmark is that target identity must be resolved at the level at which docking was performed. Multicomponent PDB structures frequently contain receptors, G proteins, signaling partners, or other cofactors. Propagating every gene in such a PDB to every chain can create artificial target aliases and inflate apparent target recovery. The final pipeline therefore uses chain-specific mapping, permits PDB-level fallback only for single-gene structures, rejects ambiguous candidate identities, and calculates GO support only from the candidate target’s own gene. This mapping discipline is necessary whenever structure-level docking results are compared with gene-level interaction benchmarks.

### 4.6 Limitations

- Molecular docking scores are approximations of physical binding and remain sensitive to protein preparation, conformational state, protonation, scoring-function limitations, and absence of cellular context [3,5].
- The structural target universe is biased toward proteins with experimentally determined structures. Using the screened structural universe as the enrichment background mitigates but does not eliminate this bias [9,10].
- GO annotations are incomplete, hierarchical, correlated, and unevenly distributed among genes [7,15,16]. Binary treatment of selected terms does not capture all semantic relationships.
- The Yamanishi benchmark contains known drug-target relationships and is not a complete record of every biologically relevant target for every compound. Apparent false positives may include unannotated or secondary interactions.
- Held-out fine-mechanistic groups are small (typically n = 2-7), limiting power and making the large raw PERMANOVA R-squared in that analysis unstable. The nonsignificant PERMANOVA p-value is therefore more informative than the raw R-squared alone.
- A formal PERMDISP analysis should be added before final submission to test whether significant PERMANOVA effects could be influenced by unequal multivariate dispersion among unbalanced groups.
- The current work is computational. A second independent DTI benchmark or prospective biochemical validation of newly prioritized targets would strengthen claims of generalizability.

### 4.7 Recommended use of the method

For practical target discovery, the results support the following hierarchy: perform proteome-scale docking; select a stable GO enrichment profile; construct a GO-associated candidate set using direct and, when appropriate, ontology-propagated annotations; retain a parallel tier of exceptional docking outliers; and apply detailed structural, pathway, expression, genetic, and experimental evidence only after this biological search-space reduction. The GO layer answers which regions of the predicted targetome warrant deeper investigation, not which single protein is definitively the target.

## 5. Conclusion

Proteome-scale docking generates broad target hypotheses but leaves an experimentally unwieldy candidate space. The corrected Yamanishi benchmark shows that stable GO Biological Process enrichment extracts reproducible biological information from these docking profiles and that the strength of this information depends on biological resolution. Broad canonical held-out classes are only weakly reproduced, whereas mechanistic and frozen fine-mechanistic analyses retain partial multivariate or local-neighborhood signal.

The strongest validated result is operational: in held-out compounds, GO-associated target space is substantially enriched for known interactions and can reduce the candidate universe more efficiently than random filtering and, at stringent search-space sizes, more efficiently than docking rank alone. Direct annotation-based additive reranking remains heterogeneous and should not be treated as a universal target score. Together, these results support GO enrichment as an intermediate biological search-space reduction and prioritization layer between proteome-scale docking and detailed target-level investigation.

## AUTHOR NOTE - REMOVE BEFORE SUBMISSION

- Complete AutoDock Vina version, docking-box definition, exhaustiveness, pose count, and any remaining preparation/scoring details in Section 2.2.
- Run and report PERMDISP as a final robustness check alongside PERMANOVA, especially for the held-out mechanistic analysis.
- Consider a second independent DTI benchmark or prospective experimental validation if targeting a higher-impact methods venue.

